# Quantitative modeling of P-TEFb mediated CTD phosphorylation identifies local cooperativity

**DOI:** 10.64898/2025.11.29.690978

**Authors:** Aaron Callenbach, Domagoj Dorešić, Robert Düster, Vanessa Nakonecnij, Erika Dudkin, Matthias Geyer, Jan Hasenauer

## Abstract

Fine-tuned regulation of RNA polymerase II (Pol II) activity is essential for accurate gene expression. A key layer of this regulation involves phosphorylation of Pol II’s C-terminal domain (CTD), a repetitive heptapeptide tail that coordinates transcription and RNA-processing factors. The kinase P-TEFb plays a major role in this process, yet its precise phosphorylation mechanism remains unclear. Previous *in vitro* studies have suggested a distributive mode of action based largely on qualitative inspection of mass spectrometry data rather than quantitative analysis. Here, we use mathematical modeling of CTD phosphorylation to explore whether local context, such as neighboring phosphorylations or directional biases, affects PTEFb activity on the CTD. Our results indicate that P-TEFb acts distributively but with pronounced local cooperativity: repeats adjacent to phosphorylated sites are modified at higher rates. We find no evidence for directional bias, although the limited positional resolution of the data precludes a definitive conclusion. These results identify local context as an important factor in P-TEFb-mediated CTD phosphorylation and establish a quantitative modeling framework for dissecting multi-site modification dynamics.

## Introduction

Accurate gene expression is fundamental to cellular identity, development, and response to environmental signals. In eukaryotes, transcription by RNA polymerase II (Pol II) is tightly coordinated with co-transcriptional RNA processing. Pol II is a multi-protein complex composed of twelve subunits (RPB1-RPB12) responsible for the transcription of proteincoding genes, as well as many non-coding RNAs.^16^ Its largest subunit, RPB1, includes a repetitive tail known as the C-terminal domain (CTD), consisting of tandem heptarepeats of the seven-amino-acid sequence Tyr–Ser2–Pro–Thr–Ser5–Pro–Ser7 (YSPTSPS) (Fig. 1A). The CTD undergoes dynamic phosphorylation that coordinates the transcription process.^8^ A distinct set of transcriptional Cyclin-dependent kinases (CDKs) directly regulate Pol II by phosphorylating its CTD. These CDKs thereby promote or preclude the binding of proteins to the CTD, enabling the specific recruitment of appropriate transcription factors throughout the transcription process.^8^

**Figure 1:**
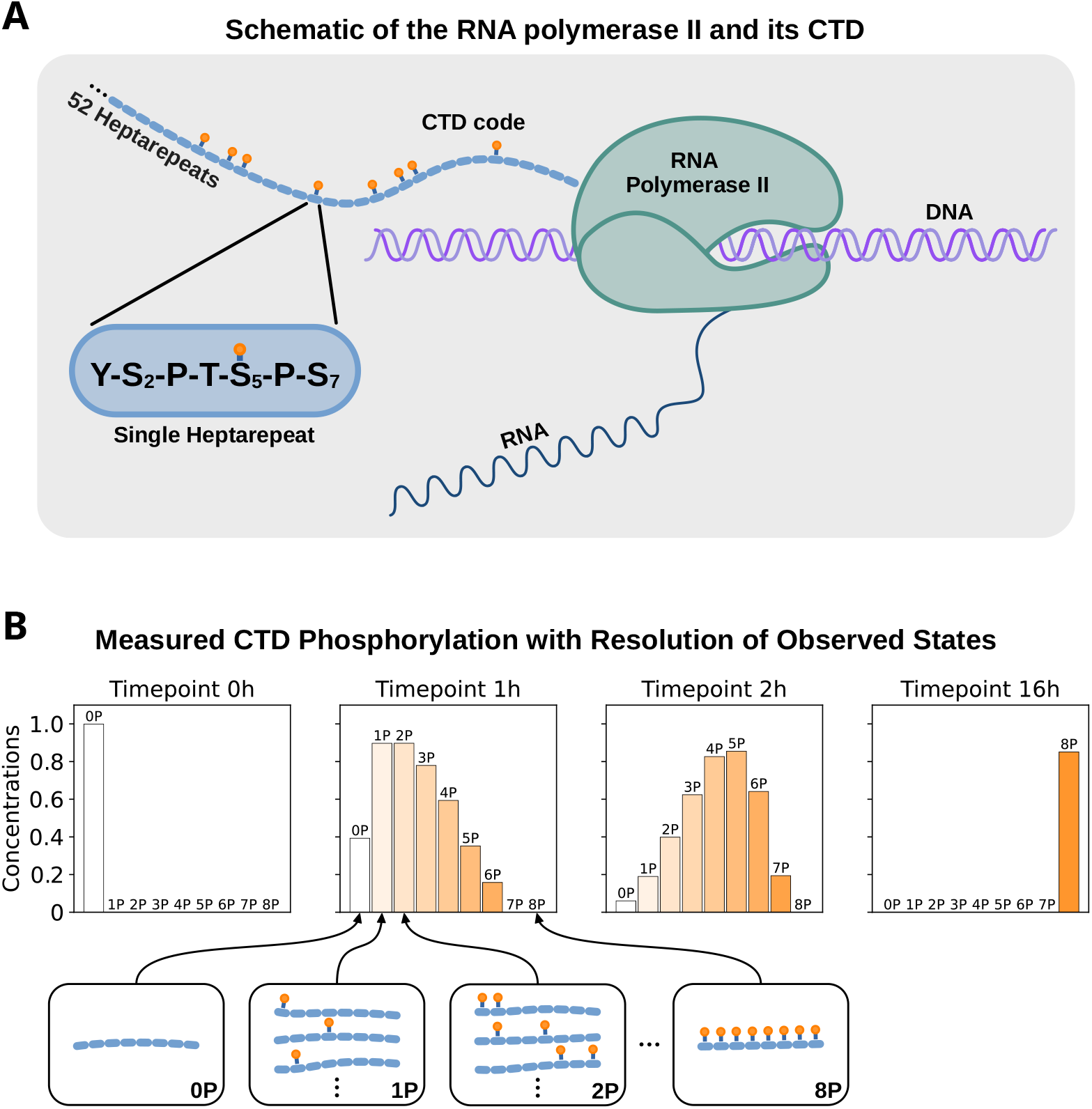
Schematic of the RNA polymerase II CTD and measured phosphorylation dynamics. (A) Schematic of the RNA polymerase II and its CTD: Blue dashes denote 52 heptarepeats of the human wild-type CTD. Orange marks represent phosphorylation of those heptarepeats. (B) CTD phosphorylation dynamics measured by mass spectrometry: mass spectrometry resolves CTD populations with 0–8 phosphorylations at different time points (0 h, 1 h, 2 h, 16 h). Bar plots show the relative abundance of phosphorylation configurations with 0 to 8 phosphorylations, with schematic CTD representations below (white-to-orange gradient indicates increasing phosphorylation).

During transcription initiation, Ser5 residues of the CTD are phosphorylated by the Kin28/CDK7 subunit of transcription factor IIH and by the Srb10/CDK8 subunit of the Mediator complex.^10,15,21,11,2^ This early modification releases Pol II from the promoter-bound preinitiation complex. ^11^ Afterwards, the CDK9 kinase of the positive transcription elongation factor b (P-TEFb) complex phosphorylates the Ser2 residues of the CTD, facilitating transcription elongation. ^21,3^ In budding yeast, the role of Ser2 phosphorylation is played by Ctk1 and Bur1, two CDK9 homologues, Ctk1 being the primary Ser2 kinase. The loss of Ctk1 nearly abolishes Ser2 phosphorylation marks.^22^ However, an *in vitro* study with human P-TEFb has revealed that it preferentially phosphorylates Ser5 over other residues.^4^ This discrepancy was addressed in a later study showing that Tyr1 phosphorylation directs the kinase activity of P-TEFb and alters its specificity from Ser5 to Ser2. ^14^ This finding highlights that the substrate specificity of P-TEFb strongly depends on the local CTD modification context. Furthermore, P-TEFb was found to be incapable of simultaneously phosphorylating Ser2 and Ser5 of the same heptarepeat,^4^ further emphasizing that the local phosphorylation state influences substrate recognition.

Beyond which residues are phosphorylated, the spatial pattern and order of CTD phosphorylation events may critically influence transcriptional regulation. Previous studies suggest that P-TEFb phosphorylates the CTD distributively rather than processively, based on time-resolved mass spectrometry distributions from hyperphosphorylation assays with human ^4^ and *Drosophila melanogaster* ^7^ P-TEFb. Yet, this conclusion was based on visual inspection and was never quantitatively confirmed, e.g. via mathematical modeling. Furthermore, it remains unexplored whether already phosphorylated sites affect the phosphorylation rate of nearby sites, either enhancing or inhibiting further modifications through altered local substrate accessibility, ^4^ or whether P-TEFb exhibits a directional preference along the CTD.

Interestingly, kinetic measurements with partially phosphorylated CTD substrates have suggested that P-TEFb may preferentially phosphorylate toward the N-terminus,^4^ hinting that existing modifications or structural features near the C-terminus could influence the direction of phosphorylation progression.

To address these questions, we formulate and compare four alternative mechanistic models: (i) fully processive phosphorylation, (ii) uniform distributive phosphorylation, (iii) distributive phosphorylation with local cooperativity (neighboring-site enhancement), and (iv) directionally biased phosphorylation. For each model, we fit the parameters with mass spectrometry time-course data^4^ and evaluate them using model selection criteria. We also assess the uncertainty and identifiability of the models’ parameters and simulations. This quantitative framework provides insight into whether P-TEFb acts processively, distributively without context dependence, or in a context-dependent manner shaped by local phosphorylation state or directional bias. Our results provide mechanistic constraints on CTD phosphorylation dynamics that inform how spatial mark patterns may regulate Pol II activity.

## Results

### Model structure and observables

To analyze CTD phosphorylation, we develop mathematical models that represent competing hypotheses about the underlying phosphorylation mechanisms. These models describe the dynamics of the abundance of CTDs with distinct phosphorylation patterns as well as the concurrent consumption of ATP.

We assess the plausibility of the model using published mass spectrometry data by Czudnochowski et al.,^4^ which quantify CTD phosphorylation configurations in an *in vitro* assay. The dataset contains 36 measurements across four time points (*t*_0_ = 0, *t*_1_ = 60, *t*_2_ = 120, and *t*_3_ = 960 minutes), reporting the distribution of CTD molecules with 0 to 8 phosphorylated heptarepeats. Each CTD consists of eight heptarepeats, and mass spectrometry resolves the number of phosphorylated repeats, but not their exact positions. Experimental evidence suggests that, under the conditions used, phosphorylation occurs selectively at either Ser2 or Ser5, but not both simultaneously within a single repeat. ^4^ Based on this, we model each heptarepeat as being either phosphorylated (1) or unphosphorylated (0), without distinguishing between the two sites.

This binary representation leads to 2^8^ = 256 possible phosphorylation configurations for heptarepeats (Fig. 2). We encode the phosphorylation configuration of a heptarepeat as vector *i* ∈ {0, 1}^8^, with the *r*-th vector element describing the state of the *r*-th heptarepeat (with 1 indicating phosphorylation). For instance, the configuration *i* = (0, 0, 1, 0, 1, 1, 0, 0) corresponds to phosphorylation at repeats 3, 5, and 6. Given this encoding, we formulate a dynamical system with for the time-dependent concentration of the different CTD forms, 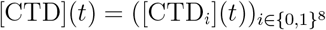, and the concentration of ATP, [ATP](*t*),

**Figure 2:**
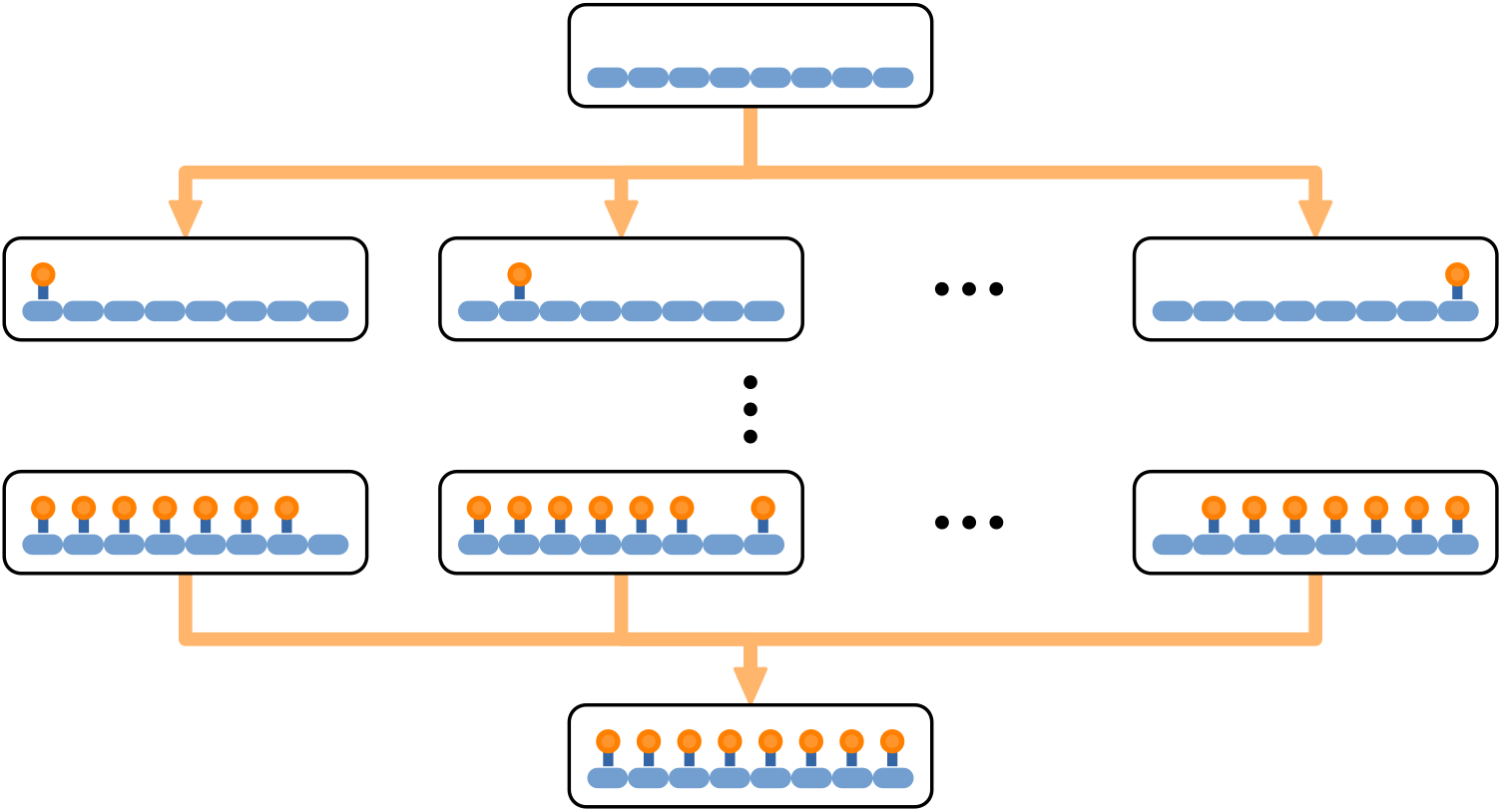
Schematic of the full model state space. Any of the eight heptarepeats (blue dashes) of the CTD molecule can be phosphorylated (orange marks) or unphosphorylated. Orange arrows indicate phosphorylation reactions.

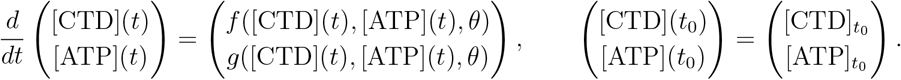

To formalize the phosphorylation dynamics, we introduce the relation *i* ≺ *j* if *i*_*r*_ ≤ *j*_*r*_ for all *r* = 1, …, 8, meaning that configuration *i* can be phosphorylated into configuration *j*. If additionally ∥*j* − *i*∥_1_ = 1, then *j* differs from *i* by exactly one additional phosphorylation. Each phosphorylation step is modeled as a reaction that converts a CTD molecule from configuration *i* to configuration *j* through the addition of one phosphate group, consuming one ATP molecule in the process. For example, the reaction

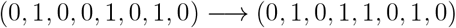

describes the phosphorylation of the fourth repeat. More generally, for all pairs *i, j* ∈ {0, 1}^8^ with *i* ≺ *j* and ∥*j* − *i*∥_1_ = 1, we include the reaction

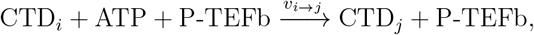

where *v*_*i*→*j*_ is the rate of phosphorylation. The kinase P-TEFb is assumed to act catalytically and is not consumed during the reaction, so its concentration [P-TEFb] remains constant over time. Furthermore, the vector field *f* follows standard mass action kinetics for all models. Specifically, the vector field for an index *i* ∈ {0, 1}^8^ is given by:

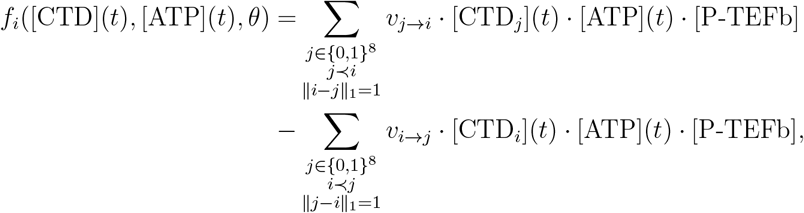

where the first sum captures the inflow from less phosphorylated configurations, while the second sum represents the outflow to more phosphorylated configurations. Additionally, the function *g* for the time evolution of ATP is given by:

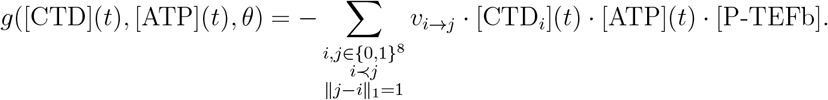

The four model hypotheses differ in the way phosphorylation rates *v*_*j*→*i*_ are defined. For example, in the uniform model, all rates are equal to the base phosphorylation rate *v*_*j*→*i*_(*θ*) = *k*_*p*_.

In addition to the base phosphorylation rate, the model includes several observational parameters required to link the model to the experimental data. Because mass spectrometry reports relative signal intensities rather than absolute molecule concentrations, we incorporate scaling parameters into the model to ensure comparability between predictions and measurements. The full parameter vector is given by:

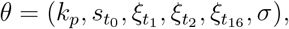

which captures both dynamical and observational aspects of the system. As already mentioned, the parameter *k*_*p*_ represents the base phosphorylation rate of P-TEFb under a context-independent uniform assumption. The parameter 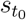 acts as a reference scaling factor, mapping the state variables of the model to the observed intensities at the initial measurement (*t*_0_). Then, to account for the systematic variation in measurement at subsequent time points, we introduce time point-specific factors 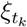, defining the effective scaling at each time point as 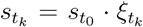. The standard deviation at each time point is also scaled using these factors, resulting in time point specific standard deviations 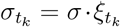 that quantify the measurement noise. In more complex hypotheses, we extend this parameter vector by including local-context sensitivity factors, such as neighbour or directional enhancement terms.

The observable mapping which relates predicted concentrations of CTDs with exactly *ℓ* (*ℓ* = 0, …, 8) phosphorylated repeats to the mass spectrometry measurements for a time point with index *k* ∈ {0, 1, 2, 3} is given by:

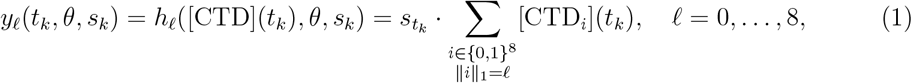

The sum is taken over all binary configurations *i* with exactly *ℓ* entries equal to one.

### Model selection points to enhanced local phosphorylation

The mechanism of CTD phosphorylation has generally been considered distributive rather than processive. To evaluate this quantitatively and to test whether local inter-repeat context further modulates phosphorylation, we fit and compare three models that encode (i) a *Fully processive* mechanism, (ii) a *Uniform distributive* mechanism, and (iii) a *Neighbouring-effect* mechanism. All of these models share the same observational model described above.

In the ***Fully processive model***, phosphorylation is initiated at one end of the CTD chain and then sequentially propagates towards the other end with a phosphorylation rate *k*_*p*_ (Fig. 3A). Processive phosphorylation could initiate at either end of the CTD chain. This would result in a reduced state space of 16 possible phosphorylation configurations. However, since the phosphorylation rate is assumed to be constant and equal for all reactions, phosphorylation started at either end would be completely symmetric. Therefore, we consider only one direction to reduce the state space even further to 9 possible phosphorylation configurations 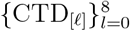, where [*ℓ*] ∈ {0, 1}^8^ denotes the multi-index for which only the first *ℓ* entries are set to 1. This reduced system results in equivalent phosphorylation dynamics to the system with 16 phosphorylation configurations. For this system with 9 possible configurations, the only non-zero kinetic rates of the model are

**Figure 3:**
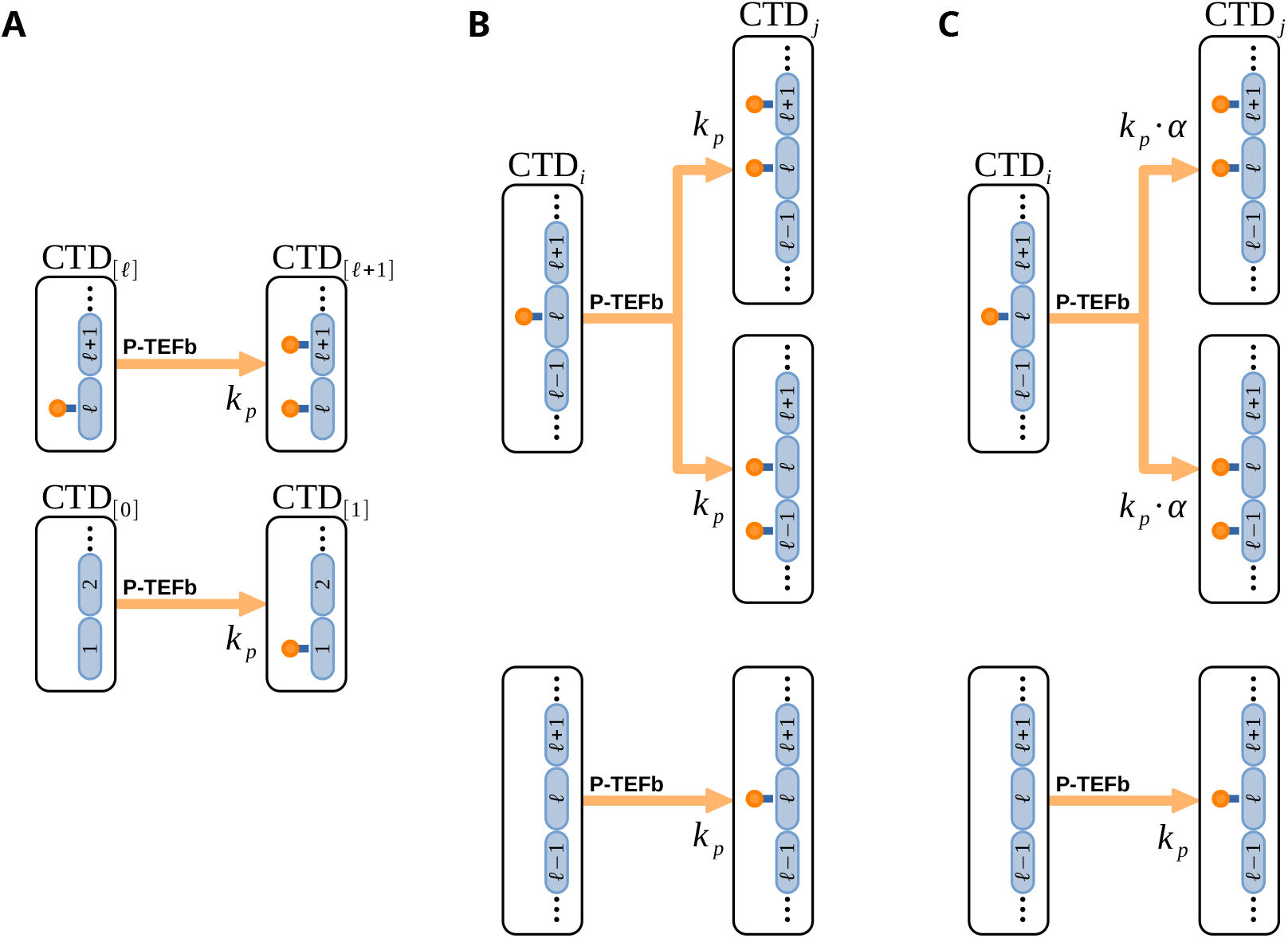
Assumptions of three model alternatives. (A) Assumptions of the *Fully processive model*: P-TEFb phosphorylates heptarepeats processively, starting from one end and continuing until the other. (B) Assumptions of the *Uniform distributive model*: P-TEFb can phosphorylate any unphosphorylated heptarepeat with identical phosphorylation rate. (C) Assumptions of the *Neighbouring-effect model*: P-TEFb exibits different phosphorylation rates for heptarepeats neighbouring already phopshorylated heptarepeats encoded by the multiplicative rate factor *α*.

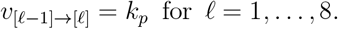

Furthermore, in a simpler case where ATP concentrations can be assumed to be approximately constant, the already-reduced system of 9 ODEs can be solved analytically. This is possible since these 9 state variables are connected in a chain of phosphorylation reactions, each state variable [CTD]_[*ℓ*]_ with one inflow and one outflow reaction. Thus, the system of ODEs can be solved inductively without the need for numerical simulation (see Materials and Methods).

In the ***Uniform distributive model***, any unphosphorylated repeat can be modified independently at the same rate (Fig. 3B). The reaction rate between two phosphorylation configurations *i* and *j* is equal to *k*_*p*_ if they differ in the phosphorylation status of a single heptarepeat, i.e.

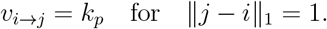

In this model, all 2^8^ = 256 phosphorylation configurations are possible. However, as all heptarepeats are phosphorylated at the same rate *k*_*p*_, any phosphorylation configuration *i* ∈ {0, 1}^8^ with *ℓ* phosphorylations can be shown to have the same probability of occurrence. Namely, for any phosphorylation configuration *i* ∈ {0, 1}^8^ the probability of randomly choosing a CTD with phosphorylation configuration *i* from the pool of all CTDs at a given time is given by:

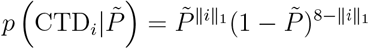

in which 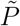 denotes the overall probability that any randomly chosen heptarepeat is phosphorylated. This probability 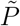 can be computed by solving an ODE system with only two state variables. Furthermore, this ODE system can be solved analytically (see Materials and Methods).

In the ***Neighbouring-effect model***, the uniform rates are scaled when the newly modified site is adjacent to an already phosphorylated repeat (Fig. 3C): if *ℓ* denotes the phosphorylated position, then

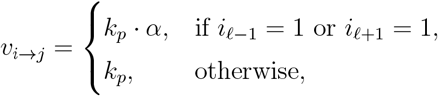

with boundary sites treated as unoccupied (*i*_0_ = *i*_9_ = 0). This breaks phosphorylation independence between sites and requires numerical integration of the full ODE systems with 2^8^ state variables.

All three models are fit to the same mass-spectrometry time course data (Fig. 1B) using identical parameter bounds (see Table S1) and multi-start optimization. Model performance is evaluated using the Negative Log-likelihood (NLLH), Akaike Information Criterion (AIC) and Bayesian Information Criterion (BIC). Convergence and runtime are inspected with waterfall plots and per-start computation times (Fig. 4).

**Figure 4:**
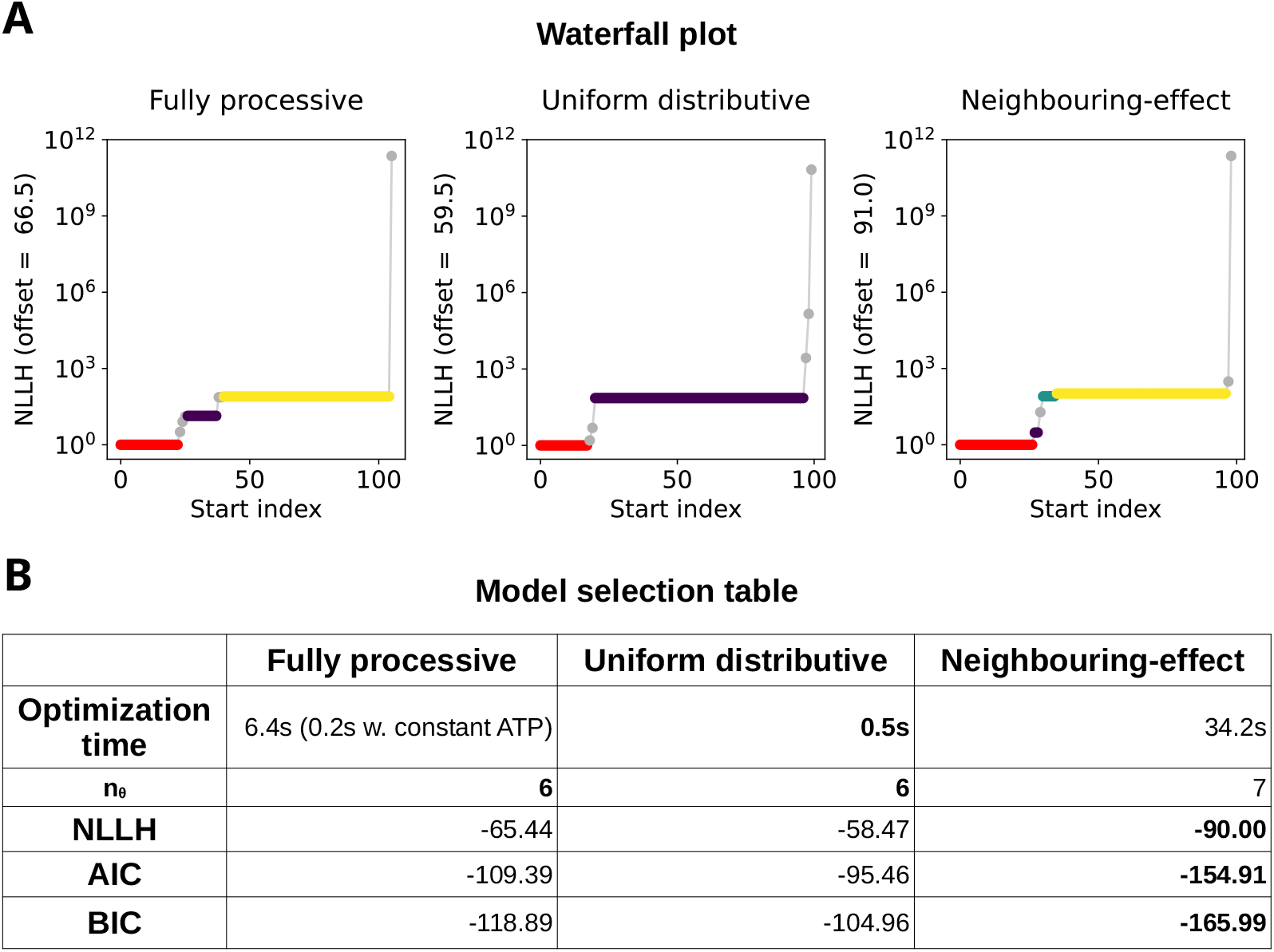
Overview of the performance of three model variants. The comparison includes the *Fully processive model*, the *Uniform distributive model*, and the *Neighbouring-effect model*. (A) The waterfall plots indicate the sorted values of the final NLLH values of all optimization starts (i.e., local optimizations). (B) Model selection table comparing optimization times, number of parameters (*n*_*θ*_), NLLH, AIC and BIC. The optimization time shows the average time to execute one local optimization of the model. A “w. constant ATP” time denotes the average computation time of a model with an assumed constant ATP amount. Lower values of the NLLH, AIC and BIC indicate better model performance. Model selection criteria (AIC and BIC) penalize increased model complexity, i.e., larger number of parameters *n*_*θ*_.

The waterfall plots indicate good convergence of optimization for all models, with more than 10% optimization starts reaching the optimal value of the objective function. Furthermore, as expected from the reduced state space of the *Fully processive model* and the analytical solution of the *Uniformly distributive model*, they are computationally ≈ 5 − 60 times faster than the *Neighbouring-effect model*. This speedup is even larger in the case of an analytical *Fully processive model* with an assumed constant ATP level. The *Neighbouring-effect model* still remains computationally feasible for the current model with 8 phosphorylation sites.

Surprisingly, the *Fully processive model* aligns better with the *in vitro* data than the *Uniform distributive model*, as reflected by lower NLLH and improved AIC/BIC scores. The *Neighbouring-effect model* further improves the fit, with a substantial reduction in both the NLLH and the model selection criteria (ΔAIC, ΔBIC *>* 10). Taken together, the results suggest that neither a purely processive nor a purely distributive mechanism can fully explain the observed *in vitro* dynamics. Hence, the *in vitro* data by Czudnochowski et al. ^4^ indicate the presence of a local phosphorylation enhancement effect.

These results motivate a closer examination of the *Neighbouring-effect model*, including parameter and state variable uncertainties. Subsequently, we explore whether the inclusion of directional preference of phosphorylation further improves model performance.

### *Neighbouring-effect model* provides a good data description with low uncertainty

Compared to simpler processive and distributive models, the *Neighbouring-effect model* is preferred by all model selection criteria. However, to judge whether the model is an adequate mechanistic description of the underlying phosphorylation mechanism, it is important to examine the adequacy of the fit (including residual diagnostics) and quantify the uncertainty in parameter estimates and phosphorylation dynamics.

The *Neighbouring-effect model* reproduces the time-course data across phosphorylation counts (0P–8P) with close agreement between simulations and observations (Fig. 5A). The model captures the early depletion of 0P and the transient accumulation of CTD concentrations of intermediate phosphorylation counts before complete phosphorylation of all CTDs. Consistent with this close agreement, the inferred measurement noise *σ* of the model is small.

**Figure 5:**
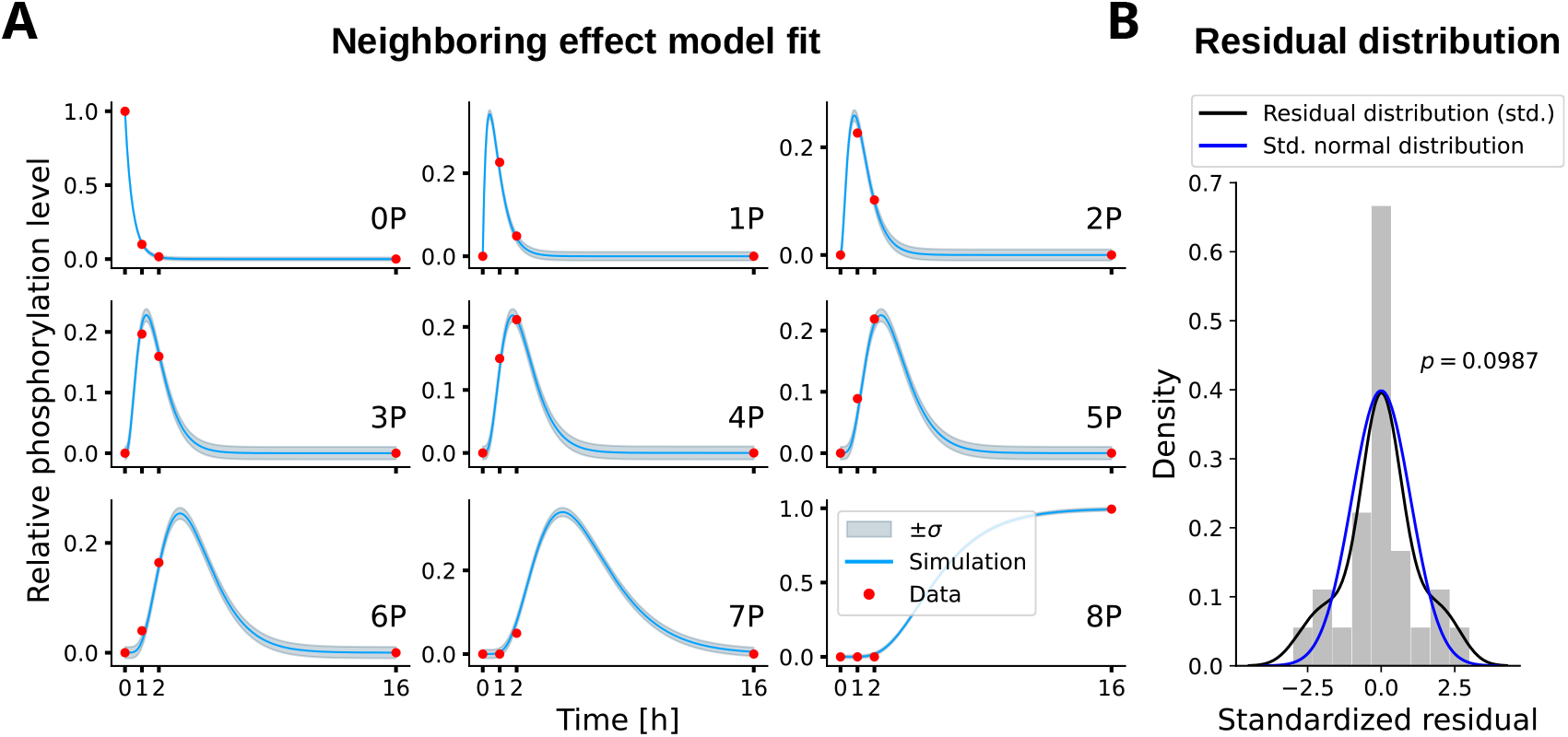
Fit of the *Neighbouring-effect model* and residual diagnostics. (A) Best-fit simulations (solid blue) with estimated *±σ* bands (gray) versus measurements (red points). The phosphorylation count is indicated with 0P-8P. (B) The histogram of the standardized residuals (gray bars) and its empirical KDE distribution (black line) is compared to the standard normal distribution (blue line).

To assess the assumption of an additive normal noise model, we compare the standardized residuals (*y*_obs_ − *y*_sim_)*/σ* and their empirical KDE distribution to the standard normal distribution 𝒩 (0, 1) (Fig. 5B). The residuals are approximately symmetric and centered at zero. Furthermore, a Shapiro-Wilk test returns *p* = 0.0987. Hence, the residuals appear to be approximately normally distribution.

To explore how well the data identifies all kinetic and observational model parameters, we compute profile likelihoods of all parameters (Fig. 6A). Each parameter profile is uni-modal and crosses the likelihood-ratio threshold on both sides of the optimum, yielding narrow confidence intervals. This is evidence of practical identifiability and low parameter uncertainty.

**Figure 6:**
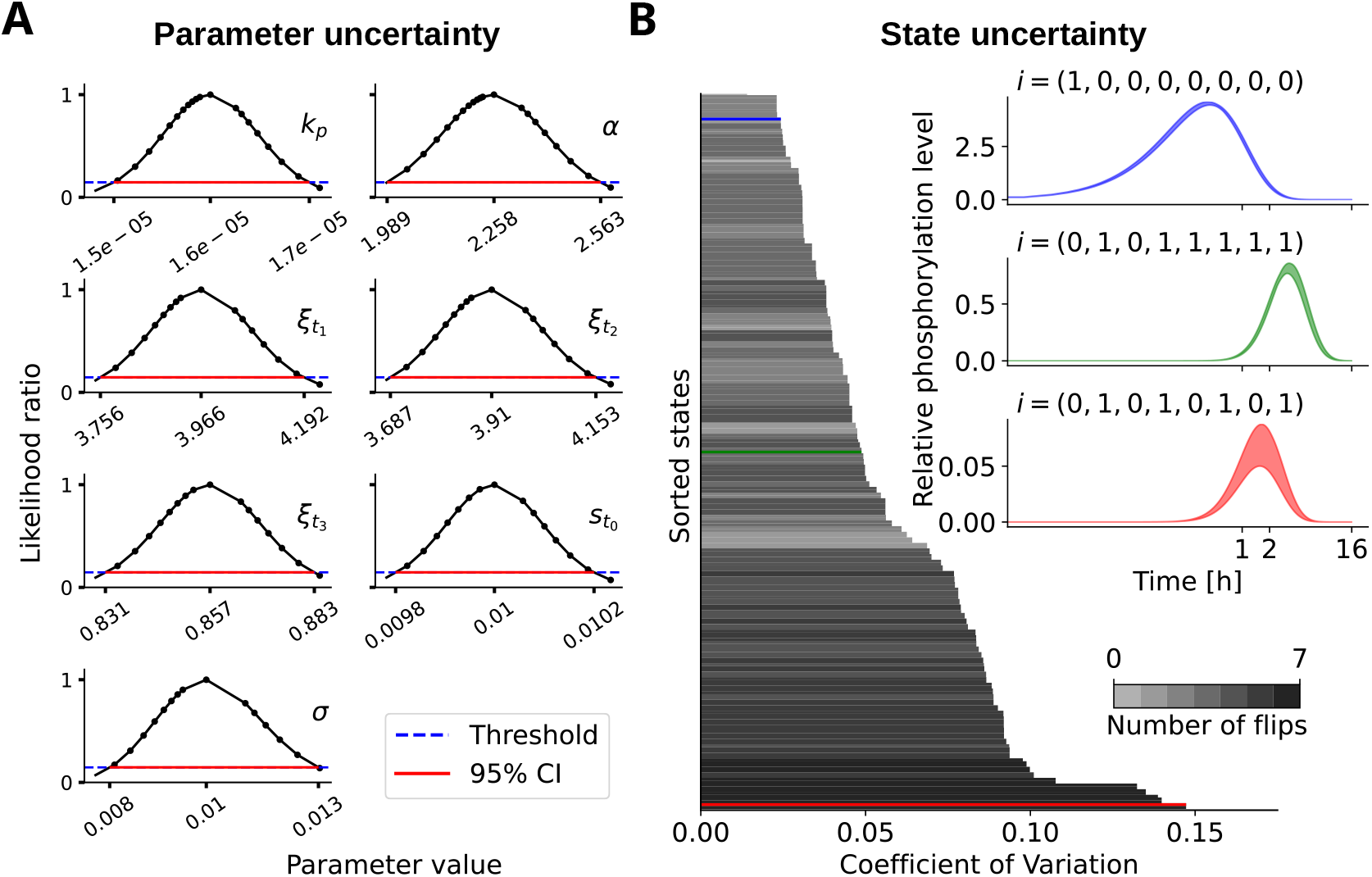
Parameter profiles and state variable uncertainty of the *Neighbouring-effect model*. (A) Likelihood-ratio profiles (black curves) for kinetic and observation parameters (*k*_*p*_, *α, ξ*_*t*1_, *ξ*_*t*2_, *ξ*_*t*16_, *s*_*t*0_, *σ*) with the likelihood-ratio threshold (blue dashed line) and 95% confidence intervals (red lines). (B) Uncertainty of the 256 phosphorylation configurations summarized by the coefficient of variation. The variability of all state variables is sorted and shown in gray bars. Phosphorylation configurations with a more alternating pattern (more 0-1 and 1-0 flips) are shown in darker gray, and simpler configurations with less alternation are show in lighter gray. To connect the coefficient of variation to uncertainty of the state trajectory, we present three examples with lower (blue), medium (green), and larger (red) coefficient of variation.

Notably, the neighbouring-effect parameter *α* is tightly bounded and its confidence interval remains above 1, indicating a confident local enhancement, rather than a reduction, of phosphorylation adjacent to already-phosphorylated sites. The *Fully processive model* and the *Uniform distributive model* give rise to equally narrow and uni-modal parameter profiles (see Fig. S1 and Fig. S2).

Lastly, we inspect how parameter uncertainty propagates through model simulation and how it affects the uncertainty in unobserved model state variables. Therefore, we collect an ensemble of parameter vectors obtained by uniformly sampling parameter values within the profile-derived confidence intervals and retaining only parameter vectors that fall inside the joint confidence region. For each parameter vector in this ensemble, we simulate the model, yielding information about the uncertainty of the state trajectories. The computation of the time-aggregated coefficient of variation for each of the 256 phosphorylation configurations,

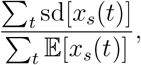

indicates an overall low uncertainty of the model state variables (Fig. 6B). As expected, states with simple phosphorylation configurations, i.e., very low phosphorylation or near-complete phosphorylation, show least uncertainty. In contrast, states with alternating phosphorylation patterns with many 0-1 and 1-0 flips, e.g., (0,1,0,1,0,1,0,1), show greater uncertainty. This is due to a stronger effect of the local enhancement factor *α* on the dynamics of these state variables. So, the uncertainty of the factor *α* is propagated into the uncertainty of these states. However, even in these cases, the uncertainty remains moderate as can be seen in the state trajectories: the uncertainty bands remain narrow relative to the trajectory peaks (Fig. 6B).

In conclusion, the *Neighbouring-effect model* provides a good quantitative description of the data with low estimated measurement noise, identifiable parameters, and low uncertainty of unobserved model state variables.

### Data does not provide evidence of a directional phosphorylation bias

In the previous section, we established the *Neighbouring-effect model* as a good description of the experimental data. However, it is not clear whether the data also support a directional preference for phosphorylation along the CTD. To test this hypothesis, we extend the *Neighbouring-effect model* by allowing for asymmetric local enhancement of phosphorylation (Fig. 7A): an upward factor *α*_up_, and a downward factor *α*_down_. Furthermore, for completeness, we include a dual factor *α*_dual_ when the target phosphorylation site is flanked by phosphorylated neighbours on both sides. Formally, for a transition *i* → *j* that phosphorylates site *ℓ*,

**Figure 7:**
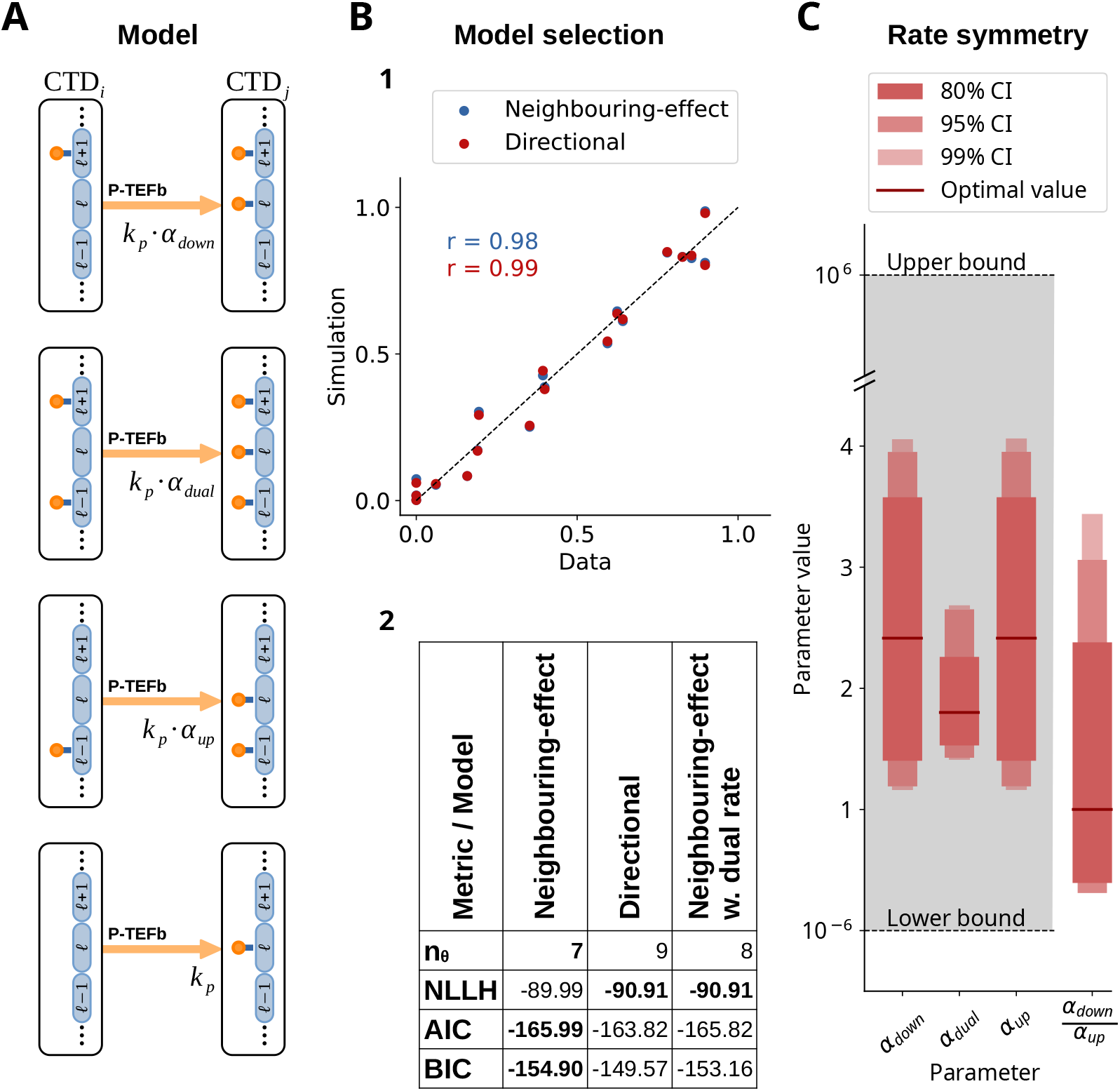
The *Directional model* and comparison with the *Neighbouring-effect model*. (A) Assumptions of the *Directional model*: schematic of base (*k*_*p*_), upward (*k*_*p*_*α*_up_), downward (*k*_*p*_*α*_down_), and dual (*k*_*p*_*α*_dual_) phosphorylation events. (B1) Data–simulation scatter plot comparing the fit of the *Neighbouring-effect model* and the *Directional model*. (B2) Values of the NLLH, AIC, BIC, and parameter counts for the *Neighbouring-effect model*, the *Directional model*, and the *Neighbouring-effect model* with a dual rate. (C) Profile-derived confidence intervals and optima of the *Directional model* for *α*_up_, *α*_down_, *α*_dual_, and the ratio *α*_down_*α*_up_.

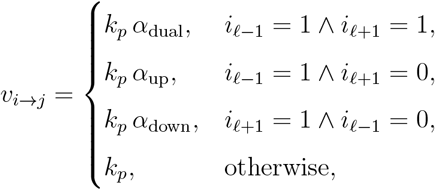

with boundary sites treated as unoccupied (*i*_0_ = *i*_9_ = 0). The observational model and estimation procedure are identical to the previous analysis.

The predicted versus observed values overlap almost perfectly for the *Neighbouring-effect model* and the *Directional model*, with data–simulation correlations *r* = 0.98 and *r* = 0.99, respectively (Fig. 7B1). This indicates that introducing separate upward/downward rates does not change the quality of the fit to the experimental data. The *Directional model* achieves a slightly lower NLLH than the *Neighbouring-effect model*, but the improvement is insufficient to compensate for the additional parameters; AIC and BIC are higher for the *Directional model* than for the *Neighbouring-effect model* (Fig. 7B2). Adding only the dual rate to the *Neighbouring-effect model* (i.e., without up/down asymmetry) achieves the same NLLH as the full *Directional model* but still worsens AIC/BIC because of the extra parameter. This equality is expected: the fitting of the *Directional model* yields 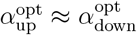 (Fig. 7C), so the asymmetry is unused and the likelihood gain stems from the dual context alone.

As for other model parameters, the profile-derived confidence intervals for *α*_up_, *α*_down_, and *α*_dual_ are narrow, indicating good practical identifiability (see Fig. S3 for full parameter profiles). Since the optimal values for *α*_up_ and *α*_down_ are essentially equal, their ratio *α*_down_*/α*_up_ is tightly concentrated around the optimal value of ≈ 1. Interestingly, *α*_dual_ is estimated below both single-neighbour factors, suggesting that a site flanked by two phosphorylated neighbours is not modified faster than with a single neighbour under these conditions (Fig. 7C).

In conclusion, within the resolution of the current *in vitro* data, there is no compelling evidence for a directional bias in CTD phosphorylation. The *Neighbouring-effect model* remains optimal according to model selection criteria.

## Discussion

CTD phosphorylation patterns regulate the timing and coordination of transcriptional processes, essential for proper gene expression and cellular function. However, the mechanistic principles that govern the kinase-mediated phosphorylation of CTD sites and establish these patterns are not fully understood. Although previous studies conclude that P-TEFb acts distributively based on visual inspection of mass spectrometry data,^4,7^ it remains unclear whether the local phosphorylation context further shapes the phosphorylation dynamics. Here we find, through a quantitative model comparison, a strong indication that P-TEFb phosphorylates the CTD distributively with local cooperativity. Repeats adjacent to already-phosphorylated sites are modified at substantially higher rates. In contrast, we find no evidence for a directional preference of P-TEFb phosphorylation. However, this may be due to the resolution of the *in vitro* data^4^ that we use for model estimation. Subsequent analysis of time-series data with individual site resolution or using pre-phosphorylated CTDs would be necessary to conclusively rule out directional bias in P-TEFb phosphorylation.

Context-dependent modulation of CTD phosphorylation has been observed in several forms, but direct evidence for inter-repeat cooperativity has been lacking. Our analysis identifies such inter-repeat effects, where phosphorylation of one heptarepeat accelerates modification of neighboring heptarepeats. Previous studies have mainly described intra-heptad dependencies. For example, pre-phosphorylation of Ser7 increases the activity of P-TEFb towards Ser2 within the same repeat. ^4^ In *Saccharomyces cerevisiae*, phosphorylation of Ser5 by Kin28 promotes the recruitment and Ser2 kinase activity of the Bur1/Bur2 complex early during transcription elongation. ^17^ However, not all CTD-modifying enzymes exhibit this dependence on the local context. For instance, O-GlcNAc transferase (OGT) catalyzes the attachment of O-linked N-acetylglucosamine residues to the Ser and Thr sites of the CTD through a purely distributive mechanism.^13^ Each modification event occurs independently of neighboring sites, producing a heterogeneous mixture of glycoforms. This contrasts with the locally cooperative phosphorylation behavior we observe for P-TEFb, highlighting that distributive modification of the CTD can arise with or without spatial interdependence between sites.

Our inference of local cooperativity implies that, even under a distributive mechanism, phosphorylation would tend to accumulate in short stretches of neighboring repeats, forming clusters of phosphorylated sites. However, this conclusion is based on *in vitro* time-course data with resolution limited to the total number of phosphorylated repeats rather than their specific positions. Consequently, the presence and extent of such clustering in vivo remain open questions and should be further explored using approaches capable of resolving CTD modification patterns at or near the level of individual repeats over time. Clear observation of transient, neighbor-to-neighbor enrichment would provide stronger evidence for inter-heptad cooperativity.

The current model represents a deliberately simplified *in vitro* setting, designed to isolate the kinetic mechanism of P-TEFb phosphorylation on a short synthetic CTD substrate. *In vivo*, CTD phosphorylation involves multiple kinases and phosphatases acting in a temporally coordinated manner throughout the transcription cycle, and such interactions are not captured here. Extending this modeling framework to include additional kinases would represent a natural next step. However, even within the *in vitro* context, expanding the model to longer CTDs introduces a severe computational challenge: the number of possible phosphorylation configurations increases exponentially with the number of repeats (2^*n*^). For the purely distributive and processive mechanisms, we derive model reductions that avoid this combinatorial explosion, but analogous reductions for the *Neighbouring-effect model* remain nontrivial. Developing such reduced or analytical formulations would be key to scaling the approach to realistic CTD lengths. More broadly, this modeling framework could be adapted to study other multisite modification systems, such as histone or RNA tail phosphorylation, where local context may similarly shape phosphorylation dynamics.

In summary, our analysis reveals that P-TEFb phosphorylates the CTD distributively but with strong local cooperativity, such that phosphorylation of one repeat enhances modification of its neighbors. This finding highlights that local context can significantly influence the dynamics of CTD phosphorylation. The model developed here offers a minimal, yet extensible framework for dissecting multisite modification dynamics and can serve as a foundation for future studies incorporating additional kinases, regulatory interactions, or longer CTD constructs.

## Materials and methods

### Experimental setup and measurement

We analyzed phosphorylation data from the *in vitro* experiments of Czudnochowski et al., ^4^ which examined how P-TEFb phosphorylates the RNAP II CTD. Mass spectrometry (ESIMS) assays were performed to determine the distribution of phosphorylation configurations on the CTD peptides. For early time points (1 and 2 hours, in addition to the 0-hour baseline), a synthetic peptide comprising eight CTD repeats (CTD[8], without a GST tag) was used as substrate. At the final 16-hour time point, a GST-tagged eight-repeat CTD (GST–CTD[8]) was employed to improve ionization efficiency. In both cases, the CTD peptide concentration was 100 *µ*M, incubated with 0.1 *µ*M P-TEFb and 3 mM non-radioactive ATP in kinase buffer (50 mM HEPES pH 7.6, 34 mM KCl, 7 mM MgCl_2_, 2.5 mM DTT, 5 mM *β*-glycerol phosphate, 0.5 mM Na_3_VO_4_) at 30^°^C. ESI-MS readouts provided relative abundances of CTD molecules with phosphorylation counts ranging from 0 to 8.

### Analytical solution for the *Fully processive model*

For the *Fully processive model*, the model output and thus also the observable functions can be calculated analytically if we assume that the ATP concentration remains constant (e.g., when ATP concentration is vastly higher than substrate concentration). This relies on the fact that the phosphorylation mechanism proceeds in a consecutive manner: Starting from one end of the CTD chain, a repeat can be phosphorylated only once the previous repeat has already been phosphorylated. As a consequence, only nine distinct phosphorylation patterns are possible, namely

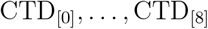

which are arranged in a simple chain of reactions. Thus, for *ℓ* ∈ {0, …, 8}, the total concentration of CTD chains with *ℓ* phosphorylated repeats is given by [CTD_[*ℓ*]_]. In addition, since the kinase is not consumed in these reactions, [P-TEFb] merely acts as a constant scalar to the reaction rate *k*_*p*_. For this reason, we will omit this scalar going forward as it can be absorbed in *k*_*p*_.

The resulting ODE system can be written in the form:

- For the unphosphorylated configuration CTD_[0]_,

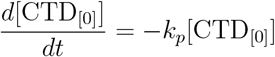
- For the intermediate configuratons CTD_[*ℓ*]_ with 1 ≤ *ℓ* ≤ 7,

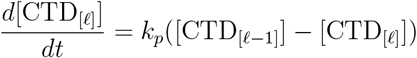
- For the fully phosphorylated configuration CTD_[8]_,

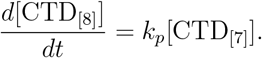

As can be seen in the data, at the initial time point all CTD chains are completely unphosphorylated, i.e.

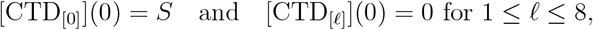

where *S* denotes the total substrate concentration. The cascadian structure of these ODEs allows us to calculate the solutions inductively: we begin with the completely unphosphorylated configuration whose ODE represents a simple exponential decay. Thus, its solution is given by:

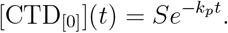

Next, we assume that for an *ℓ* ∈ {1, …, 7} the following has already been shown for the intermediate configuration CTD_[*ℓ*−1]_:

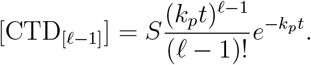

The equation for [CTD_[*ℓ*]_] takes the form

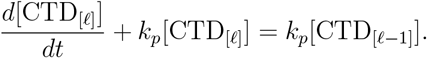

Applying the induction hypothesis and multiplying by 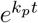 yields

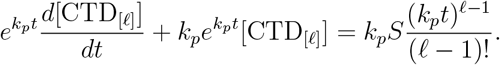

By the reversed product rule this is equivalent to

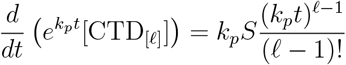

and hence

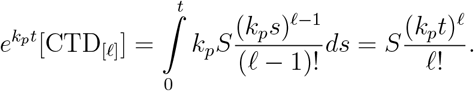

Thus, each intermediate observable (1 ≤ *ℓ* ≤ 7) takes the form of a scaled Poisson distribution term

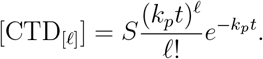

For the fully phosphorylated configuration, we do not need to solve its differential equation explicitly. Instead, we can make use of the conservation of substrate. This is possible since the total amount of substrate in the system remains the same as every CTD chain must be in exactly one phosphorylation configuration and there is no degradation or synthesis taking place. Thus,

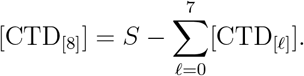

Together, the observable mapping can be calculated as

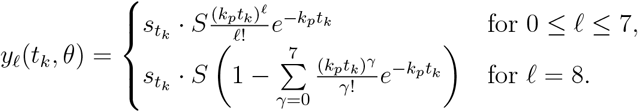

The appearance of Poisson terms in the intermediate configurations is not accidental. In fact, the phosphorylation mechanism in the *Fully processive model* can be understood directly as a Poisson counting process: At each intermediate step, only a single phosphorylation event is possible, and each occurs with the same constant rate *k*_*p*_. This setup mirrors precisely the definition of a Poisson process where events happen one at a time, independently and with a fixed rate. Let *N* (*t*) be a Poisson counting process with rate *k*_*p*_. Then the probability that exactly *ℓ* events have occurred by time *t* is

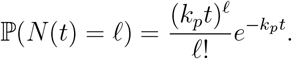

Compared with our solutions, we see that the observables are simply scaled versions of these probabilities,

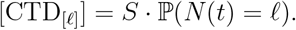

However, this Poisson interpretation only holds when the concentration of ATP is approximately constant. Once ATP dynamically changes, the effective reaction rate 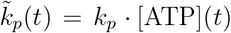 is no longer constant because it depends on the varying ATP concentration. Since a defining property of a Poisson process is that events occur independently and with a constant rate, this time dependence violates the Poisson assumption. Consequently, when ATP dynamics are included, the simple analytical solution derived above no longer applies, and the system must be solved numerically.

### Analytical solution for the *Uniform distributive model*

Since the *Uniform distributive model* assumes no cross-interactions between CTD repeats and all repeats follow identical kinetics, the total concentration of phosphorylated repeats across all CTD chains can be represented by a single reaction:

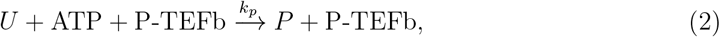

where *U* and *P* denote the total concentrations of unphosphorylated and phosphorylated repeats, respectively. To ensure these variables represent total repeat concentrations, the initial value of *U* is set to the total substrate concentration scaled by the number of repeats per CTD chain, i.e., *U*_0_ = 8 · *S*. Additionally, the total substrate concentration is conserved over time, so we can represent the concentration of the unphosphorylated repeats as *U* = *U*_0_ − *P*. Thus, reaction (2) gives rise to the following ODE system with a conserved quantity:

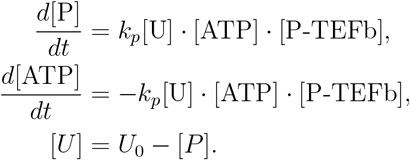

Furthermore, with *Ũ*= [*U*]*/U*_0_ we represent the fraction of unphosphorylated repeats and with 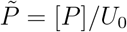 the fraction of phosphorylated repeats. Consequently, these fractions satisfy 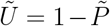. To solve the system analytically, we first note that 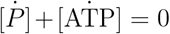, which implies that

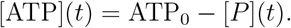

As before, we assume that the constant factor [P-TEFb] is absorbed into *k*_*p*_. Substituting the algebraic equation for [ATP] into the first ODE yields,

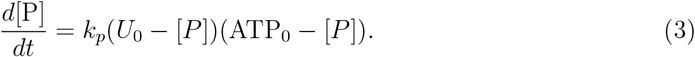

When ATP_0_ = *U*_0_, equation (3) simplifies to

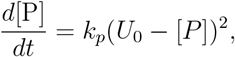

which can be solved by separation of variables to obtain

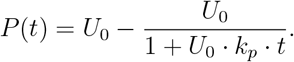

For ATP_0_ ≠ *U*_0_, we rewrite equation (3) as

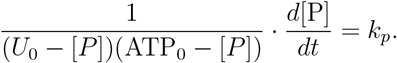

Using partial fraction decomposition, the left term becomes

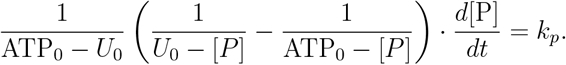

Integrating both sides and applying the initial condition [*P*](0) = 0 yields

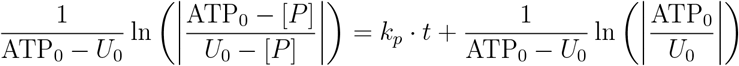

Solving for *P* gives

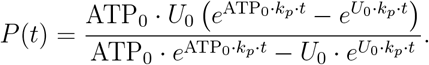

Combining both cases, we obtain the following closed-form solution:

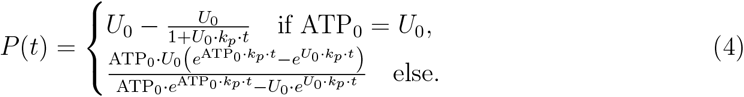

Now, to determine the temporal evolution of the distribution of CTD configurations, we first note that there are no cross-interactions between repeats and all phosphorylation events occur with identical kinetics. Therefore, the phosphorylation configurations of individual repeats are statistically independent. Consequently, the concentration of CTD chains exhibiting a specific phosphorylation pattern *i* ∈ {0, 1}^8^ can be expressed as the product of the fractions associated with each repeat being in its respective phosphorylation configuration, scaled by the total substrate concentration. Hence,

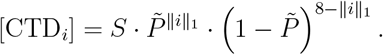

For the parameter estimation, the concentration of specific phosphorylation patterns is not relevant but only the overall concentration of CTDs with a specific number of phosphorylations, which is given by:

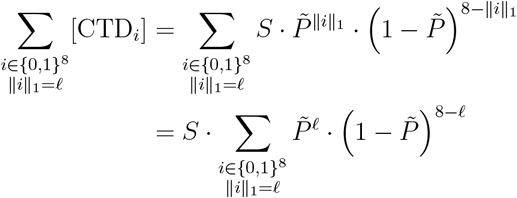

Since choosing a phosphorylation pattern with *ℓ* phosphorylated repeats amounts to selecting *ℓ* out of 8 sites, the total number of such configurations is 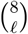. Thus, the concentration of CTD chains with exactly *ℓ* phosphorylated repeats follows a binomial distribution with parameters *n* = 8 and time-dependent success probability 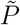, scaled by *S*:

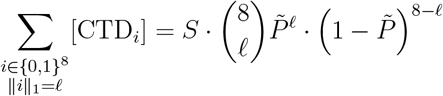

This shows that, instead of explicitly modeling all 2^8^ possible phosphorylation configurations, it is sufficient to track the total phosphorylation dynamics. The full distribution of CTD configurations can then be directly inferred via this binomial relationship. The model observables can then be written as

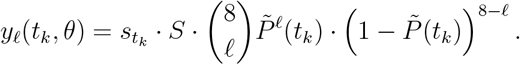

Together with the analytical solution for *P*, and thus 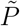, the observables can be calculated entirely analytically.

### Numerical model simulation

Time-course simulations of the ODE models were performed using the CVODES solver from the SUNDIALS suite, through AMICI’s interface.^6^ CVODES provides efficient and accurate integration of stiff systems using adaptive step-size and order control, ensuring numerical stability across a wide range of model dynamics. Simulations were performed with a relative tolerance of 10^−7^ and an absolute tolerance of 10^−16^.

### Model parameterization

All models in this manuscript are parameterized using maximum likelihood estimation. We assume additive and normally distributed measurement noise. Thus, the relation of the model observables (1) with exactly *ℓ* (= 0, …, 8) phosphorylated repeats at a time point with index *k* ∈ {0, 1, 2, 3} to the respective observed mass spectrometry data point 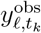 is given by:

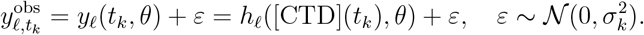

The conditional probability density of observing a specific data point at a specific time point given the model observable *y*_*ℓ*_(*t*_*k*_, *θ, s*_*k*_) and noise parameter *σ*_*k*_ is given by:

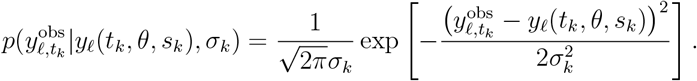

We assume all measurements are mutually independent, so the likelihood function is given as a product of conditional probabilities:

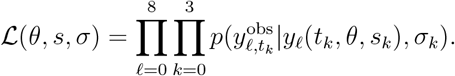

As commonly done, instead of maximizing the likelihood function, for better numerical stability, we minimize the negative log-likelihood that is given by:

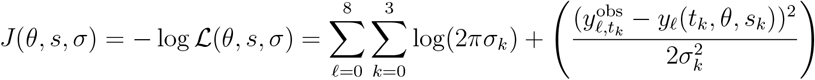

We denote the maximum likelihood estimate as the minimum of this objective function

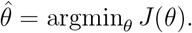

We minimize the objective function using multi-start minimization with 128 local minimizations per model. For this, we employ gradient-based optimization using the trust region optimizer Fides. ^5^ Gradients of the objective function are computed analytically for analytical models and computed via adjoint sensitivity analysis using AMICI ^6^ for all other models. Adjoint sensitivity analysis was performed with a relative tolerance of 10^−7^ and an absolute tolerance of 10^−16^.

### Model selection criteria

To compare the performance of different models in a principled way, we employ the **Akaike Information Criterion** (AIC)^1^ and the **Bayesian Information Criterion** (BIC).^20^ Both measures aim to balance goodness-of-fit with model complexity, penalizing models that use more parameters to prevent overfitting. AIC is defined as

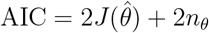

where *n*_*θ*_ denotes the number of parameters in the model and *J* the negative log-likelihood. Therefore, models that achieve a good fit (low NLLH) will be rewarded while excessive model complexity (high *n*_*θ*_) will be penalized. Thus, in general a lower AIC value indicates a more favorable trade-off between model fit and model complexity. BIC extends this idea by also incorporating the number of data points *n*_data_:

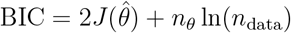

### Uncertainty quantification and profile likelihoods

We express the uncertainty of parameter estimates using their respective confidence intervals. We compute these intervals using the profile likelihood approach based on the likelihood-ratio test. The likelihood-ratio test for a parameter vector *θ* is defined via the test statistic:

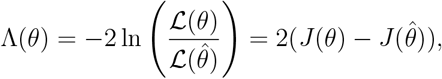

where ℒ denotes the likelihood function. In the asymptotic case of a large number of data points, the distribution of Λ(*θ*) can be approximated by the chi-square distribution 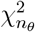 with degrees of freedom equal to the number of parameters.^23^ The confidence region of significance level *α* is defined as

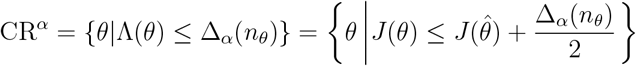

where Δ_*α*_(*n*_*θ*_) denotes the *α*-quantile of the 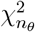 distribution. For a single parameter, the profile likelihood is defined as

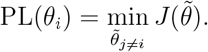

Similarly to the confidence region, the profile likelihood-based confidence interval is defined as

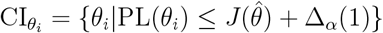

The profile likelihood provides not only a numerical estimate of parameter uncertainty through the confidence interval width, but also a visual and qualitative assessment of how sensitively the likelihood responds to changes in each parameter. Furthermore, we assess practical identifiability using profile-derived confidence intervals: a combination of model and data is said to be practically identifiable if the confidence intervals of all parameters are finite.

### Implementation and availability

The model definition and parameter estimation tasks were formulated using the PEtab (v0.5.0) format. ^19^ Models were created with PySB (v1.16.0) ^12^ and encoded in SBML.^9^ Parameter estimation and uncertainty analysis were performed using pyPESTO (v0.5.5). ^18^ The model simulations used AMICI (v0.31.2),^6^ and the trust region optimizer Fides (v0.7.8) ^5^ was used for numerical optimization. All SBML and PEtab files, and the code and instructions to perform all analyses, are available at https://zenodo.org/records/17736181.

### Declaration of usage of AI tools in the writing process

Portions of the text were refined with the assistance of ChatGPT, which was used for language editing and improving clarity. The authors reviewed and approved all generated text and are responsible for the final content.

## Funding

This work was supported by the Deutsche Forschungsgemeinschaft (DFG, German Research Foundation) under Germany’s Excellence Strategy (EXC 2047—390685813, EXC 2151—390873048), by the European Union by ERC grant INTEGRATE (GA number 101126146), and by the University of Bonn (via the Schlegel Professorship of J.H.). M.G. is supported by the European Research Council (ERC Advanced Grant NalpACT).

## Author Contributions

J.H., and M.G. conceptualized the project and acquired financial support. A.C., V.N., R.D., D.D., E.D., M.G. and J.H. developed the mathematical model. A.C., D.D., and J.H. calibrated and analyzed the model. J.H., A.C., and D.D. wrote the initial draft and prepared the visualization. All authors discussed the results and commented on the manuscript.

## Supporting information

**Figure S1:**
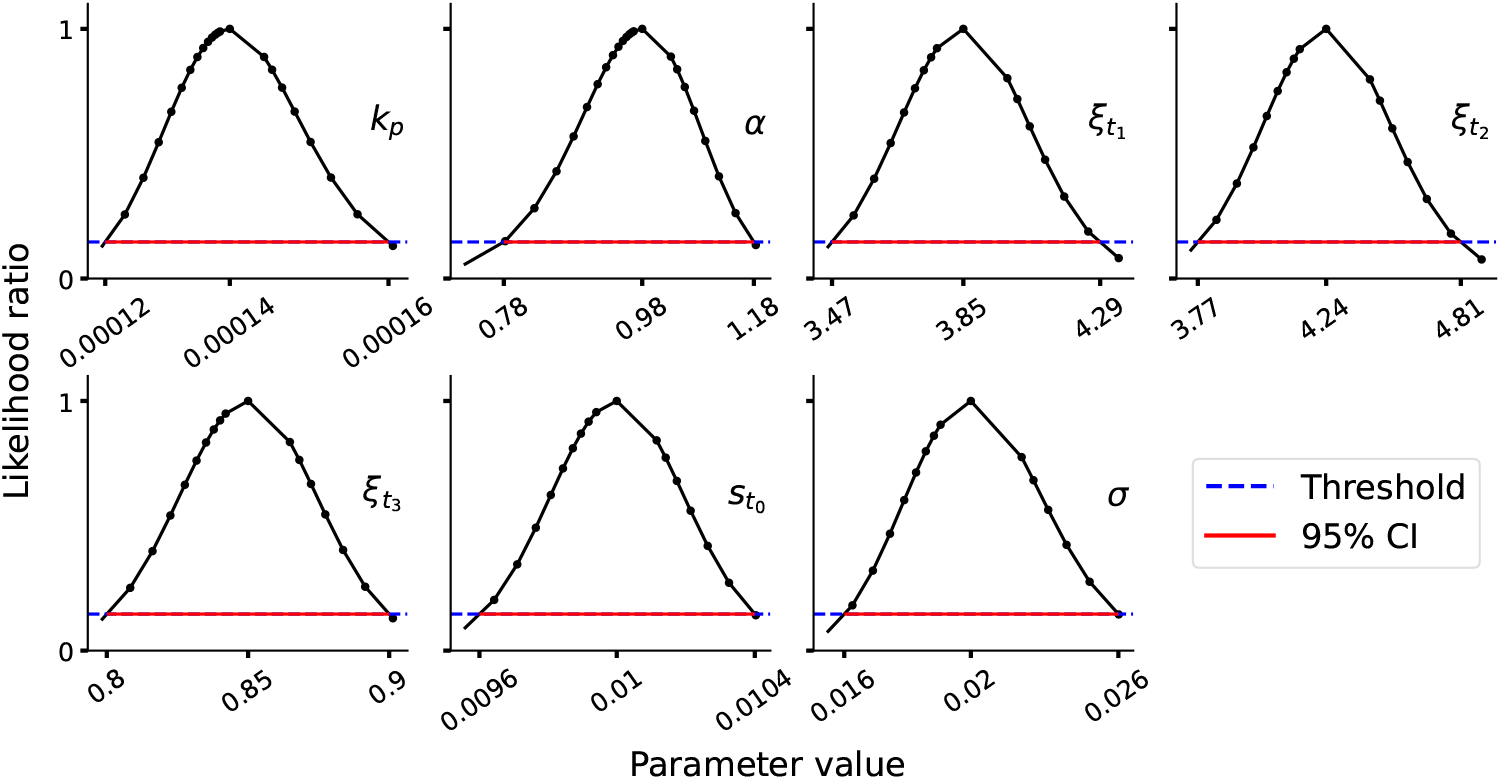
Profile likelihoods for parameters of the *Fully processive model*. Black curves show the obtained profile likelihood (maximum obtained likelihood for the fixed parameter value) normalized by the maximum likelihood from optimization. The red line marks the 95% confidence threshold, and the blue dashed line indicates the CI cutoff.

**Table S1:**
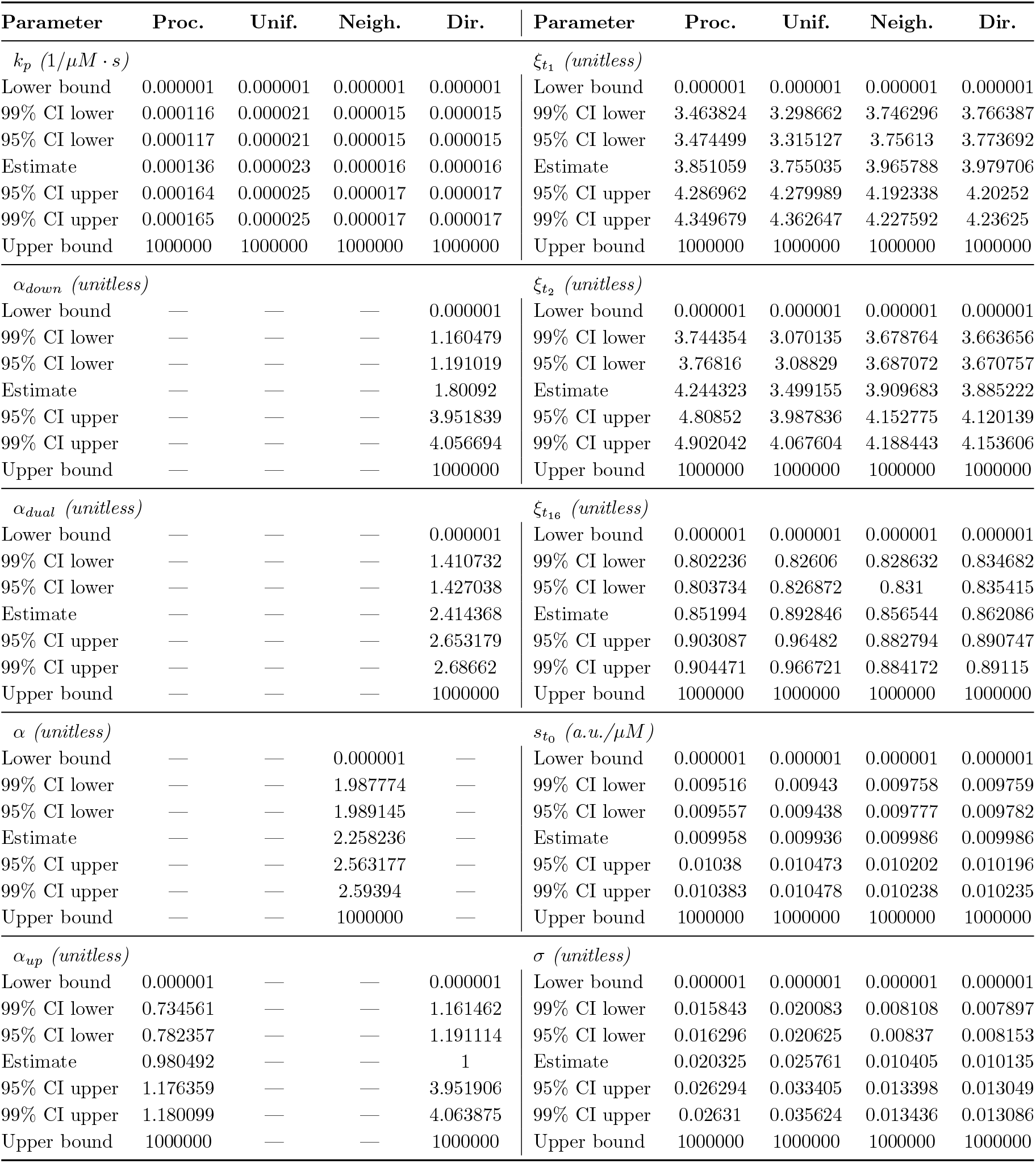
Parameters of all models. Contains all parameters used in all models with their parameter bounds, confidence intervals across models, and parameter units.

| Parameter | Proc. | Unif. | Neigh. | Dir. | Parameter | Proc. | Unif. | Neigh. | Dir. |
| --- | --- | --- | --- | --- | --- | --- | --- | --- | --- |
| $k_p$ ($1/\mu M \cdot s$) | | | | | $\xi_{t_1}$ (*unitless*) | | | | |
| Lower bound | 0.000001 | 0.000001 | 0.000001 | 0.000001 | Lower bound | 0.000001 | 0.000001 | 0.000001 | 0.000001 |
| 99% CI lower | 0.000116 | 0.000021 | 0.000015 | 0.000015 | 99% CI lower | 3.463824 | 3.298662 | 3.746296 | 3.766387 |
| 95% CI lower | 0.000117 | 0.000021 | 0.000015 | 0.000015 | 95% CI lower | 3.474499 | 3.315127 | 3.75613 | 3.773692 |
| Estimate | 0.000136 | 0.000023 | 0.000016 | 0.000016 | Estimate | 3.851059 | 3.755035 | 3.965788 | 3.979706 |
| 95% CI upper | 0.000164 | 0.000025 | 0.000017 | 0.000017 | 95% CI upper | 4.286962 | 4.279989 | 4.192338 | 4.20252 |
| 99% CI upper | 0.000165 | 0.000025 | 0.000017 | 0.000017 | 99% CI upper | 4.349679 | 4.362647 | 4.227592 | 4.23625 |
| Upper bound | 1000000 | 1000000 | 1000000 | 1000000 | Upper bound | 1000000 | 1000000 | 1000000 | 1000000 |
| $\alpha_{down}$ (*unitless*) | | | | | $\xi_{t_2}$ (*unitless*) | | | | |
| Lower bound | — | — | — | 0.000001 | Lower bound | 0.000001 | 0.000001 | 0.000001 | 0.000001 |
| 99% CI lower | — | — | — | 1.160479 | 99% CI lower | 3.744354 | 3.070135 | 3.678764 | 3.663656 |
| 95% CI lower | — | — | — | 1.191019 | 95% CI lower | 3.76816 | 3.08829 | 3.687072 | 3.670757 |
| Estimate | — | — | — | 1.80092 | Estimate | 4.244323 | 3.499155 | 3.909683 | 3.885222 |
| 95% CI upper | — | — | — | 3.951839 | 95% CI upper | 4.80852 | 3.987836 | 4.152775 | 4.120139 |
| 99% CI upper | — | — | — | 4.056694 | 99% CI upper | 4.902042 | 4.067604 | 4.188443 | 4.153606 |
| Upper bound | — | — | — | 1000000 | Upper bound | 1000000 | 1000000 | 1000000 | 1000000 |
| $\alpha_{dual}$ (*unitless*) | | | | | $\xi_{t_{16}}$ (*unitless*) | | | | |
| Lower bound | — | — | — | 0.000001 | Lower bound | 0.000001 | 0.000001 | 0.000001 | 0.000001 |
| 99% CI lower | — | — | — | 1.410732 | 99% CI lower | 0.802236 | 0.82606 | 0.828632 | 0.834682 |
| 95% CI lower | — | — | — | 1.427038 | 95% CI lower | 0.803734 | 0.826872 | 0.831 | 0.835415 |
| Estimate | — | — | — | 2.414368 | Estimate | 0.851994 | 0.892846 | 0.856544 | 0.862086 |
| 95% CI upper | — | — | — | 2.653179 | 95% CI upper | 0.903087 | 0.96482 | 0.882794 | 0.890747 |
| 99% CI upper | — | — | — | 2.68662 | 99% CI upper | 0.904471 | 0.966721 | 0.884172 | 0.89115 |
| Upper bound | — | — | — | 1000000 | Upper bound | 1000000 | 1000000 | 1000000 | 1000000 |
| $\alpha$ (*unitless*) | | | | | $s_{t_0}$ (*a.u./μM*) | | | | |
| Lower bound | — | — | 0.000001 | — | Lower bound | 0.000001 | 0.000001 | 0.000001 | 0.000001 |
| 99% CI lower | — | — | 1.987774 | — | 99% CI lower | 0.009516 | 0.00943 | 0.009758 | 0.009759 |
| 95% CI lower | — | — | 1.989145 | — | 95% CI lower | 0.009557 | 0.009438 | 0.009777 | 0.009782 |
| Estimate | — | — | 2.258236 | — | Estimate | 0.009958 | 0.009936 | 0.009986 | 0.009986 |
| 95% CI upper | — | — | 2.563177 | — | 95% CI upper | 0.01038 | 0.010473 | 0.010202 | 0.010196 |
| 99% CI upper | — | — | 2.59394 | — | 99% CI upper | 0.010383 | 0.010478 | 0.010238 | 0.010235 |
| Upper bound | — | — | 1000000 | — | Upper bound | 1000000 | 1000000 | 1000000 | 1000000 |
| $\alpha_{up}$ (*unitless*) | | | | | $\sigma$ (*unitless*) | | | | |
| Lower bound | 0.000001 | — | — | 0.000001 | Lower bound | 0.000001 | 0.000001 | 0.000001 | 0.000001 |
| 99% CI lower | 0.734561 | — | — | 1.161462 | 99% CI lower | 0.015843 | 0.020083 | 0.008108 | 0.007897 |
| 95% CI lower | 0.782357 | — | — | 1.191114 | 95% CI lower | 0.016296 | 0.020625 | 0.00837 | 0.008153 |
| Estimate | 0.980492 | — | — | 1 | Estimate | 0.020325 | 0.025761 | 0.010405 | 0.010135 |
| 95% CI upper | 1.176359 | — | — | 3.951906 | 95% CI upper | 0.026294 | 0.033405 | 0.013398 | 0.013049 |
| 99% CI upper | 1.180099 | — | — | 4.063875 | 99% CI upper | 0.02631 | 0.035624 | 0.013436 | 0.013086 |
| Upper bound | 1000000 | — | — | 1000000 | Upper bound | 1000000 | 1000000 | 1000000 | 1000000 |

**Figure S2:**
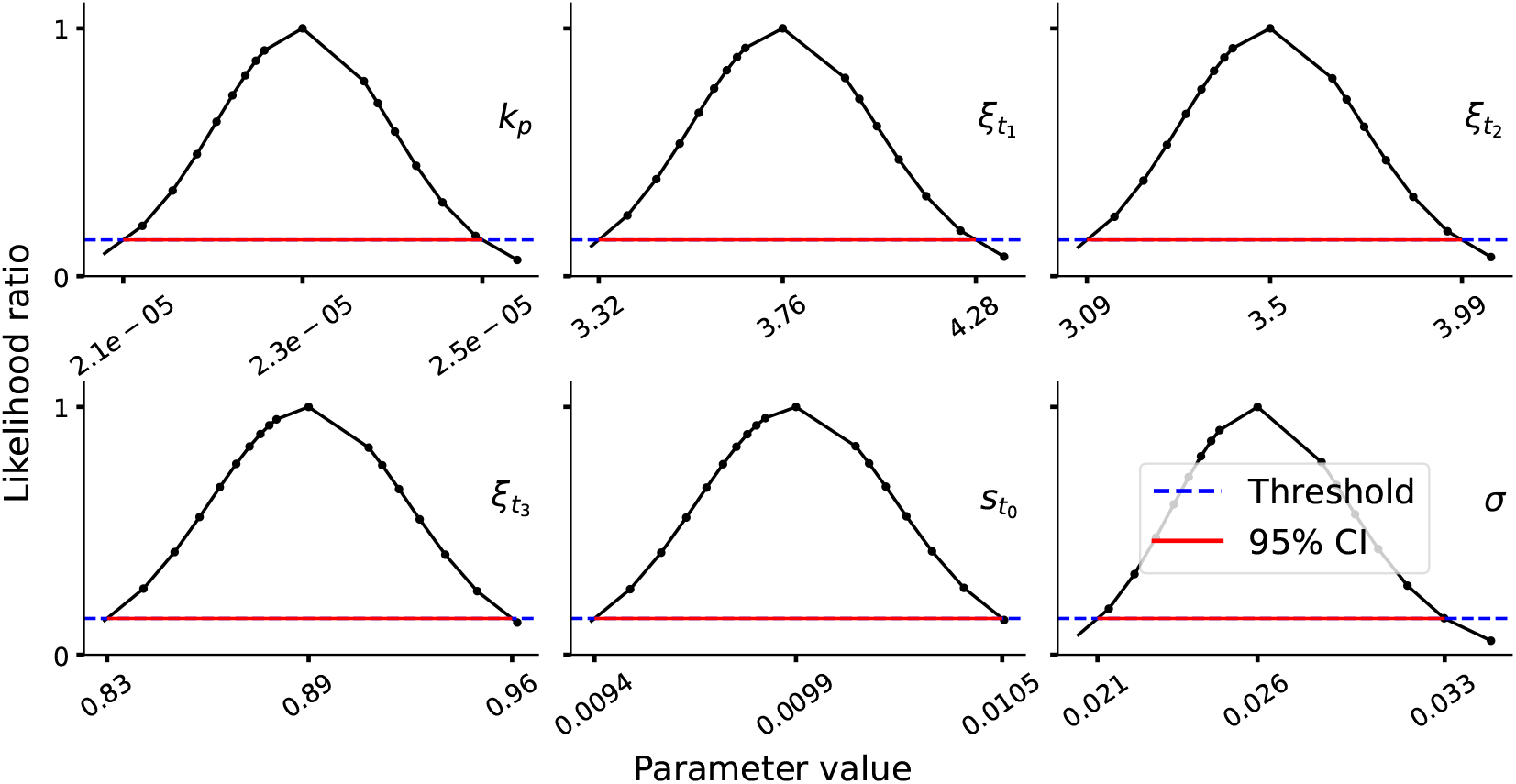
Profile likelihoods for parameters of the *Uniform distributive model*. Black curves show the obtained profile likelihood (maximum obtained likelihood for the fixed parameter value) normalized by the maximum likelihood from optimization. The red line marks the 95% confidence threshold, and the blue dashed line indicates the CI cutoff.

**Figure S3:**
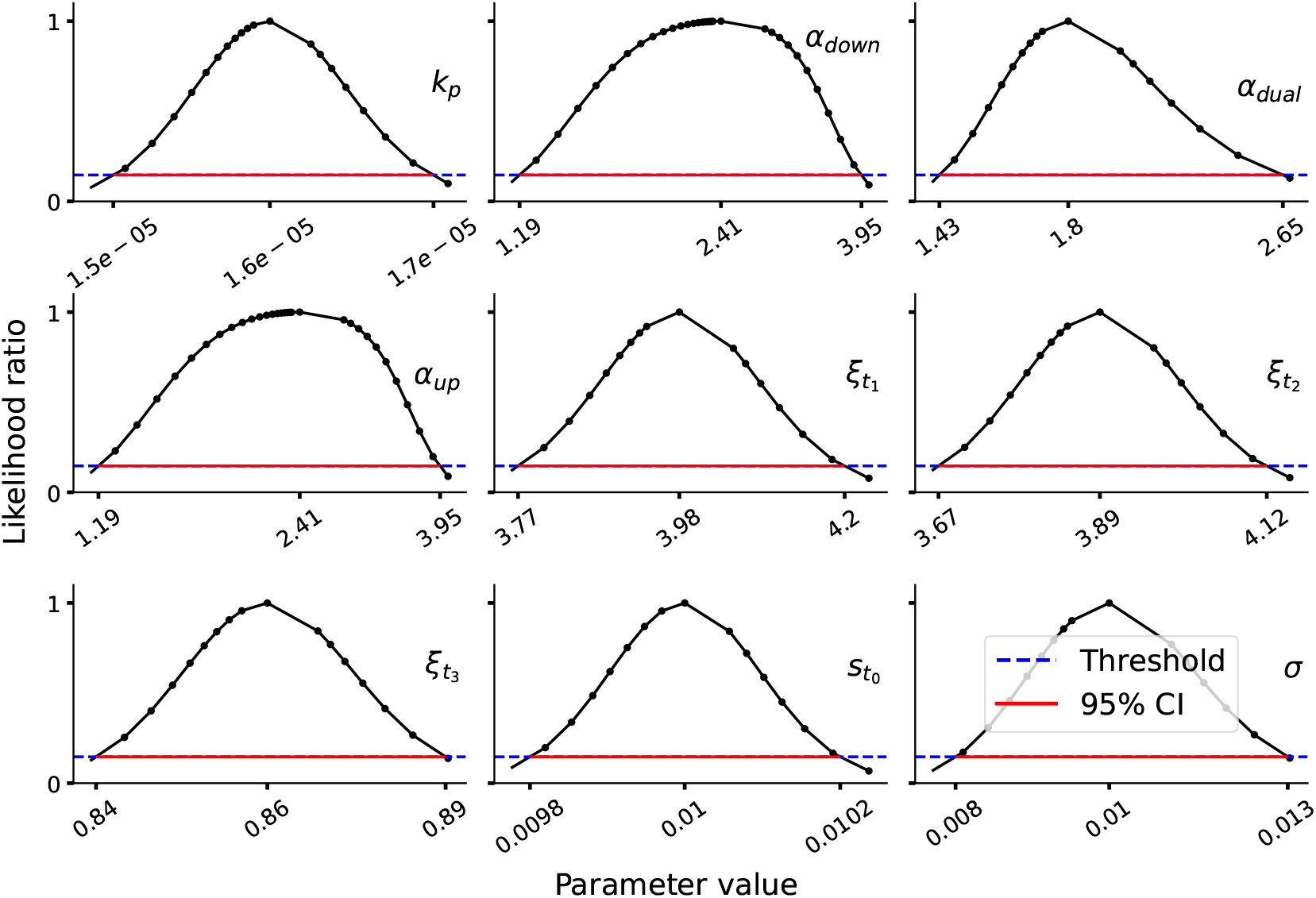
Profile likelihoods for parameters of the *Directional model*. Black curves show the obtained profile likelihood (maximum obtained likelihood for the fixed parameter value) normalized by the maximum likelihood from optimization. The red line marks the 95% confidence threshold, and the blue dashed line indicates the CI cutoff.

## Notes

https://zenodo.org/records/17736181

### Competing Interest Statement

The authors have declared no competing interest.

